# Carbon catabolite repression in pectin digestion by phytopathogen *Dickeya dadantii*

**DOI:** 10.1101/2021.04.04.438409

**Authors:** B. Shiny Martis, Michel Droux, William Nasser, Sylvie Reverchon, Sam Meyer

**Affiliations:** Université de Lyon, INSA Lyon, Université Lyon 1, CNRS UMR 5240, Laboratoire de Microbiologie, Adaptation et Pathogénie, 11 avenue Jean Capelle, 69621 Villeurbanne, France

## Abstract

The catabolism of pectin from the plant cell walls plays a crucial role in the virulence of the phytopathogen *Dickeya dadantii*. In particular, the timely expression of *pel* genes encoding major pectate lyases is essential to circumvent the plant defense systems and induce a massive pectinolytic activity during the maceration phase. While previous studies identified the role of a positive feedback loop specific to the pectin degradation pathway, here we show that the *pel* expression pattern is controlled by a metabolic switch between glucose and pectin. We develop a dynamical and quantitative regulatory model of this process integrating the two main regulators CRP and KdgR related to these two sources of carbon, and reproducing the concentration profiles of the associated metabolites, cAMP and KDG respectively, quantified using a new HPLC method. The model involves only 5 adjustable parameters, and recapitulates the dynamics of these metabolic pathways during bacterial growth together with the regulatory events occurring at the promoters of two major *pel* genes, *pelE* and *pelD*. It highlights their activity as an instance of carbon catabolite repression occurring at the transcriptional regulatory level, and directly related to the virulence of *D. dadantii*. The model also shows that quantitative differences in the binding properties of common regulators at these two promoters resulted in a qualitative different role of *pelD* and *pelE* in the metabolic switch, and also likely in conditions of infection, explaining their evolutionary conservation as separate genes in this species.

## Introduction

The expression of virulence genes is known to respond to metabolic changes in many pathogenic species, including *Salmonella enterica* (1), *Vibrio cholerae* (2), *Helicobacter pylori (3), Pseudomonas aeruginosa* (4), *Escherichia coli* and *Shigella spp* (5), revealing a tight interconnection between virulence and metabolic functions (6). This link is particularly conspicuous in the phytopathogen *Dickeya dadantii*, a soft rotting Gram-negative bacterium that attacks a wide range of plant species, including many crops of economical importance (7). In a first asymptomatic phase of infection, *D. dadantii* colonizes the intercellular space (apoplast) where it grows on available simple sugars produced by the plant. The symptomatic phase, maceration, is characterized by the degradation of pectin, a complex sugar present in the plant cell walls, resulting in soft rot, the main visible symptom (8). The success of the infection process crucially depends on *D. dadantii*’s ability to switch from glucose to pectin catabolism in a timely and efficient manner, in order to overcome attacks by the plant’s defense systems (9).

The depolymerization of the pectin polysaccharide is achieved by a variety of pectinases. Among these, endo-pectate lyases (Pels) are known to play a prominent role and considered as major virulence factors (8), especially those encoded by the *pelD* and *pelE* paralogous genes (10). In presence of pectin, the expression of *pel* genes is induced via a positive feedback loop: the binding of the main repressor KdgR at their promoters is relieved in presence of its cofactor KDG, itself resulting from the degradation of pectin by Pel enzymes (11). This nonlinear mechanism has been formerly proposed to explain the strong boost in *pel* expression required for a fast switch to the maceration phase (12). However, culture experiments showed that pectin is degraded early during bacterial growth ((12) and Fig. 1), whereas *pel* expression peaks only hours later, close to the transition to stationary phase (Fig. 1D). In this paper, we propose a new model for this regulatory loop, where the expression of pel genes is triggered not by the degradation of pectin alone, but rather by the shift in metabolic uptake from glucose to pectin. We show that this shift is controlled by a carbon catabolite repression (CCR) mechanism, *i*.*e*., the selective uptake of a preferred carbon source (glucose) by repression of the catabolism of other sources (here pectin). It presents many similarities but also interesting differences with the classical glucose/lactose example in *Escherichia coli*, in particular regarding dynamical properties required for a successful infection (13).

**Figure 1.**
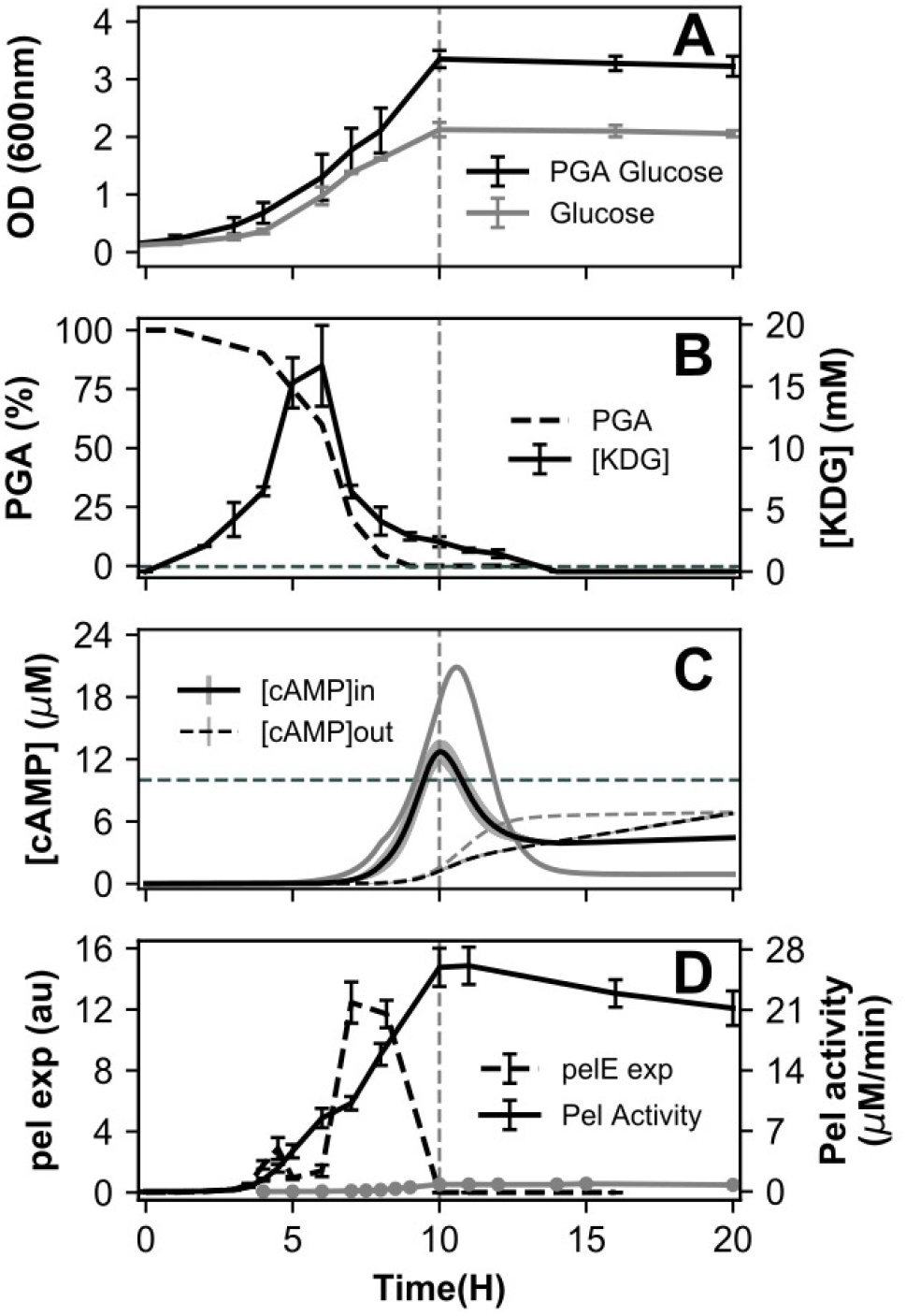
**(A)** Growth curves of *D. dadantii* in M63 minimal medium supplemented with glucose (gray) or glucose+PGA (black). **(B)** Time course of PGA degradation (dashed line, data from (12)) and intracellular KDG concentration measured by HPLC (solid line). The horizontal dashed line (0.4 mM) indicates the KDG-KdgR affinity: above this level, repression is relieved and *pel* expression is strongly induced. **(C)** Time course of measured extracellular cAMP concentration (dashed line) and inferred intracellular cAMP concentration (solid line, with shaded confidence area), for cells grown in glucose (gray) or glucose+PGA (black). The dashed horizontal line (10 µM) indicates the cAMP-CRP binding affinity (16): above this level, *pel* expression is activated. Raw data points are shown in Supplementary Figures S5 and S6. **(D)** *pelE* expression measured by qRT-PCR (dashed line) and total Pel enzymatic activity in glucose (gray) or glucose+PGA (black) media.

Although the regulation of pectin degrading *pel* genes involves the combined action of at least a dozen different transcription factors and nucleoid-associated proteins (8), the strongest of these regulators monitored in the cell are KdgR and cAMP receptor protein (CRP) (8). Since the latter is also the main catabolite activator protein controlling the glucose metabolic switch in *E. Coli* (14), we hypothesized that the combined action of these two regulators could explain the time course of *pel* expression through variations in the concentration of their respective co-factors, KDG and cAMP. To validate this hypothesis, we used a highly sensitive method to measure the concentrations of these metabolites, based on a derivatization of the compounds followed by HPLC quantification. Based on these data, we present a quantitative dynamical model of the regulation pathway. The binding affinities of both regulators with their cofactors as well as *pel* operators were formerly measured (12, 15) as well as other key parameters of the system, requiring only 5 adjustable parameters in the model. The latter explains the dynamics of Pel production during bacterial growth when both glucose and pectin are available, as well as the specific role of the recently diverged paralogous genes *pelE* and *pelD* in pectinolysis (10), explaining why this divergence could constitute an evolutionary advantage for pathogenesis.

## Results

### Pectin degradation and KDG concentration peak occur before Pel production boost

The production of pectate lyases was monitored during *D. dadantii* growth in minimal medium supplemented with glucose or glucose+polygalacturonate (PGA), a simple form of pectin (Fig. 1D). The latter acts as an inducer, boosting the production of enzymes by a factor of around 30, which was attributed to the indirect activation of *pel* genes by the metabolite KDG resulting from PGA degradation (11, 12, 15): when KDG binds the regulator KdgR, it relieves transcriptional repression by the latter, thus triggering a positive feedback loop of pectin degradation (Fig. 2). However, this scenario is challenged by the dynamics of PGA degradation (Fig. 1B, data from (12)), which occurs extremely rapidly after around 5h of growth, hours before most Pel enzymes are produced (close to transition). To explain this delay, a very slow import of the extracellular pectin degradation products (oligogalacturonides, UGA) and/or subsequent intracellular conversion into KDG inducer was invoked (Fig. 2, the detailed pathway of pectin degradation is shown in Supplementary Figure S1); yet this scenario seems unrealistic, since depolymerisation rather than import is thought to be the rate-limiting step of this pathway, and because such a slow kinetics would be a major obstacle for the fast activation of *pel* genes by pectin required for efficient plant infection (9).

**Figure 2.**
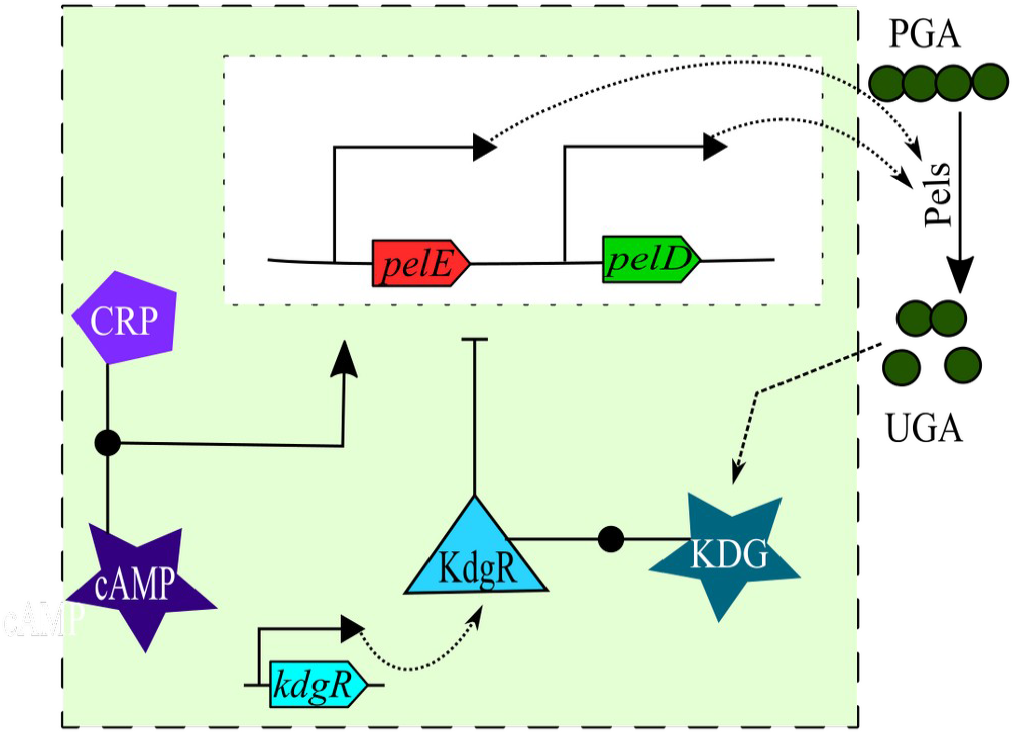
Diagram of the components and reactions considered in the model. The two metabolites, cAMP and KDG, indirectly activate the expression of *pel* genes by binding their associated regulators: cAMP allows CRP to bind and activate the promoters, whereas KDG relieves KdgR repression, respectively. Pel enzymes are instantaneously exported and degrade PGA (pectin) into UGA, which is imported and converted into KDG, triggering a positive feedback loop. KdgR is synthesized at a constant rate.

Resolving this discrepancy required a direct measurement of intracellular KDG concentration. This was achieved using a new method based on the derivatization of the carbonyl group of the molecule with o-phenylenediamine and subsequent HPLC quantification. Elution with a nonlinear increasing gradient of trifluoroacetic acid and acetonitrile allows separating KDG from its close structural analogues, as described in (17). Samples collected at regular time intervals clearly indicate that the KDG inducer peak occurs almost immediately after PGA degradation, after around 6h of growth (Fig. 1B), confirming that the import of oligosaccharides and further degradation steps into KDG are extremely fast and cannot be the limiting steps for strong *pel* expression. Most strikingly, the lifetime of KDG is also quite short in the cell, since the concentration has already dropped to half its value one hour after its peak time (the growth rate is by far insufficient to explain this decrease through a dilution effect). This profile should be compared to the expression profile of the major pectate lyase gene *pelE* measured by qRT-PCR (Fig. 1D), which peaks almost entirely after the concentration of KDG inducer has dropped. Since the cellular concentration of the KdgR repressor does not vary considerably with time (12), these observations show that the timing of *pel* expression cannot be explained by the KDG-KdgR regulatory pathway alone. The question then arises to propose an alternate or additional regulatory mechanism.

### Pel production time is mainly controlled by cAMP accumulation

*pel* genes exhibit a particularly complex regulation, probably optimized for the specific requirements of virulence where any error in their expression time and strength results in the detection, attack, and ultimately, destruction of the pathogen by the host’s defense systems (9). This regulation involves global transcription factors (CRP) and nucleoid-associated proteins (Fis, H-NS, IHF), which act in combination with DNA supercoiling (18), as well as more classical regulators of the pectin catabolic pathway (KdgR, PecT, PecS, MfbR) (8). Several of these regulators might contribute in relating *pel* expression to the metabolic state of the cell, such as the Fis repressor that is mostly present in early exponential phase, or DNA supercoiling (18). However, the latters’ quantitative effect on *pel* expression is known to be significantly milder than that of CRP and KdgR (18, 19), which are considered as the main regulators and whose binding and regulatory properties on several *pel* promoters were therefore analysed with much detail, both *in vitro* (15, 19) and *in vivo* (11, 20). We took advantage of this accumulated knowledge to develop a quantitative modeling of the regulatory dynamics of the system.

The first step consisted in quantifying the second main contribution of the regulatory signal, *i*.*e*., the concentration of the metabolite cAMP, which triggers a conformational change in CRP that allows it to bind most of its operator sites and play its activating role (Fig. 2 and (19)). This quantification was performed by HPLC after derivatization using 2-chloroacetaldehyde for fluorescence detection (21) (see Materials and Methods). Samples were collected at regular time intervals during growth, and cAMP concentrations were quantified in the extracellular medium. Internal cAMP concentrations were then inferred from these latter by means of a kinetic model of cAMP export and import, as previously proposed (22) (see Supplementary Information). The resulting intracellular cAMP concentration profile (Fig. 1C) entirely matches previous profiles obtained by both direct (23, 24) and indirect measurements in *E. coli* (22), with a very low level observed during exponential growth in glucose, a sharp peak at the transition to stationary phase, and lower but nonzero concentrations in the latter.

The cAMP concentration time course was monitored during growth on glucose with or without PGA (Fig. 1C). The profiles are qualitatively similar, except that the peak is somewhat sharper when glucose is the only sugar source present, as we expected. In both conditions, the binding of a significant fraction of CRP proteins present in the cell (the affinity is indicated as a dashed horizontal line), subsequent promoter binding by the cAMP-CRP complex and Pel enzyme production boost are thus expected to occur around the transition to stationary phase, as indeed observed (Fig. 1D). In summary, while KDG acts as an inducer of Pel production (as visible in the relative amounts of enzymes produced in the two media), the timing of this production is thus rather controlled by cAMP, in contrast with previous hypotheses (12). Note that, for the concentration of PGA used in these experiments, the levels of internal KDG concentration reached in the cell are around 40 times higher than those required to bind KdgR and relieve the repression (0.4 mM, horizontal dashed line in Fig. 1B), explaining why the inducing effect can last hours after KDG concentration has started to decrease. In other words, only a tiny fraction of Pels, produced very early, is sufficient to degrade the whole amount of pectin present in our culture medium, whereas the actual peak of Pel production occurs only hours later, when cAMP starts to accumulate in the cell close to the transition to stationary phase.

### Combined regulation of *pel* genes through KDG and cAMP metabolites

We now translate these qualitative observations into a quantitative model, schematized in Fig. 2, which explicitly incorporates two pel genes with major virulence effect, *pelD* and *pelE*. To facilitate the understanding, we first describe the model without distinguishing these two genes which share the same regulators and produce enzymes of similar activity, leaving the analysis of their differential effects to the next section.

The dynamics of all components (bacterial cell density and concentrations of enzymes, regulators and metabolites inside/outside the cells) are simulated over time, according to a set of coupled differential equations (see Supplementary Information). Most parameters were obtained from experimental knowledge (Supplementary Table S2), and the 5 remaining parameters (Pel production rates and kinetic parameters controlling KDG synthesis and degradation, see Supplementary Table S3) were numerically adjusted to reproduce the observations of Fig. 1.

The essential steps and results of the modeling are illustrated on Fig. 3. The KDG profile results from the degradation of PGA by Pel enzymes previously produced and exported (A). While KdgR is synthesized in the cell at a constant rate, it is mostly bound by KDG when the latter is present in sufficient quantity (over 0.4 mM), resulting in the unbinding of *pel* gene promoters (B) and relieve of KdgR repression (C) compared to cells grown with glucose only (J). The expression of *pel* genes is computed based on a thermodynamic model of transcription (25), with experimental values of KdgR/CRP affinities and activation factors (Table 1). To illustrate the effect of each regulatory signal, we compute activation profiles (Figs. 3 C, E, J, L), representing the gene expression fold-change due to repressor (<1) or activator (>1) binding, and proportional to the statistical weight of the unbound or bound state of the corresponding regulatory protein (here KdgR and CRP) respectively. Because the KdgR-KDG regulatory system involves a positive feedback loop of highly nonlinear behavior, reproducing the measured KDG peak is the most sensitive step in the numerical tuning procedure since it depends on several kinetic parameters of unknown value (Supplementary Table S3), whereas the regulatory part of the model entirely relies on knowledge-based parameter values.

**Figure 3.**
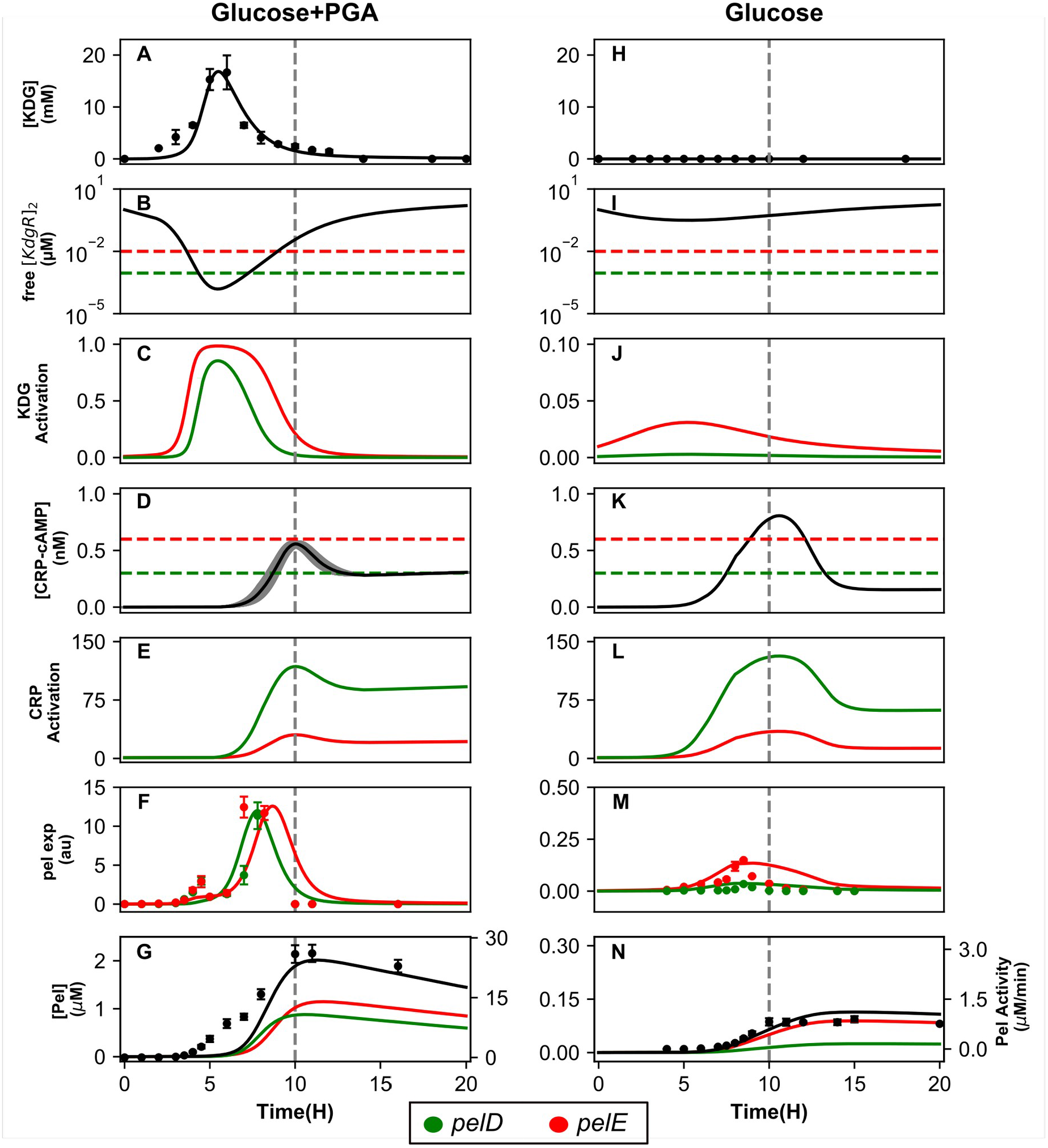
Modeling of *pel* regulatory pathways and expression in bacteria grown in minimal medium supplemented with glucose+PGA (left column, A-G) or glucose (right, H-N). Model results are shown as solid lines, and experimental data as dots. The transition time is shown as a dashed gray vertical line. **(A)** KDG intracellular concentration superimposed with HPLC direct measurement data exhibiting a peak after around 6 hours of growth; **(B)** concentration of intracellular free KdgR dimer, with horizontal dashed lines showing the binding affinities of the dimer for the *pelE* (red) and *pelD* (green) promoters, indicating the thresholds of repression; **(C)** *pelE* and *pelD* (depicted in red and green respectively throughout the figure) activation curves based on KdgR binding, exhibiting a de-repression in the middle of exponential phase due to high intracellular levels of KDG; **(D)** intracellular concentration of the cAMP-CRP complex inferred from HPLC measurement of cAMP in the medium (Fig. 1B); the gray area is the 95% confidence interval due to experimental variations, and dashed horizontal lines indicate the binding affinities for *pelD/E* promoters; **(E)** *pel* activation curves based on cAMP-CRP binding, with a boost occuring at transition; **(F)** *pel* expression time course based on combined regulation by KdgR and CRP, superimposed with qRT-PCR datapoints peaking around 8h; **(G)** extracellular concentration of Pel enzymes, either from individual genes (red and green) or the combination of both (black), superimposed with measurements of enzymatic activity (dots) reflecting total Pel enzymes, including those not considered in our modeling. **(H-N)** Same legends as A-G: in absence of PGA, the dilution of KdgR during growth is sufficient to partly relieve *pelE* repression, while cAMP induces expression of both *pels* close to transition. The timing and expression levels of both genes is accurately reproduced by the model in both conditions, as well as the inducing effect of PGA.

**Table 1.** Experimental regulatory parameter values, and expression levels measured and obtained by the model after optimization, for *pelD* and *pelE*.

| Promoter | *peID* | *peIE* |
| --- | --- | --- |
| KdgR binding affinity (nM) | 0.9 | 10 |
| cAMP-CRP binding affinity (nM) | 0.3 | 0.6 |
| cAMP-CRP activation factor | 181 | 62 |
| PGA induction factor (exp) | 330 | 83 |
| PGA induction factor (model) | 324 | 93 |

In order to keep the modeling as simple as possible, we considered the internal cAMP concentration profile as an input information (Fig. 1C). cAMP binds CRP with a measured affinity, assuming a constant concentration of the protein in the cell, resulting in a concentration peak of the cAMP-CRP complex at the transition (Fig. 3D). In turn, this complex binds and activates *pel* gene promoters based on measured affinities and activating factors (Tab. 1 and Fig. 3E) (15). The peak in *pel* expression thus results from the combination of the two latter regulatory signals, which are separated in time but exhibit a short overlap after around 7-8 hours of growth, in relatively good agreement with our qRT-PCR data both in presence and absence of pectin (Figs. 3 F and M). Similarly, the production level and timing of total Pel enzymes match the experimental profiles, with an approximate 30-fold induction by PGA (Figs. 3 G and N). Note that the measured Pel enzymatic activity exhibits a slow accumulation already at an earlier time than in the model, and matching with early secondary peaks of *pelE/D* expression after around 4h. This discrepancy might be explained by two factors: (1) the activity profile contains contributions from various *pel* genes not considered explicitly in the model and (2) additional regulators modulating *pel* expression are disregarded here (see next section).

### Quantitative differences in regulatory sequences underpin qualitatively different roles for paralogous genes

In the modeling, we explicitly considered two separate *pel* genes, *pelD* and *pelE*, which play a major role in virulence as observed from pathogenicity assays (26), and were therefore studied extensively (18, 27). These paralogous genes result from a recent duplication event from a common ancestor present in other *Dickeya* species (10), respond to the same regulators and encode proteins of similar enzymatic activity, yet were shown to play different roles during bacterial growth and plant infection (10). Since the regulatory parameters are experimentally available and differ between the two promoters due to point mutations (Tab. 1), we addressed the question, whether the modeling could help understanding these distinct roles.

The main known difference between these genes is the level of induction by pectin. Whereas they exhibit a comparable expression level in presence of PGA, *pelE* still has a significant level of expression in absence of inducer, whereas that of *pelD* is undetectable (10, 28). In the modeling, any observed difference in expression results primarily from the respective binding affinities of the KdgR repressor, which differ by a factor of ten (1 nM for *pelD*, 10 nM for *pelE* promoter). With a cellular concentration of KdgR in the micromolar range as measured *in vivo* (12), both promoters are mostly bound by the repressor in absence of PGA in the medium (Fig. 3I); however, because of the affinity difference, unbinding events are far more frequent at the *pelE* than *pelD* promoter, hence the relatively higher basal expression level of the former (Fig. 3J). Still in the glucose medium, in mid-exponential phase, fast growth results in KdgR dilution by cell divisions, thus weakening its repressive effect on *pelE* and favoring an expression peak before transition time (Fig. 3I). In the presence of PGA, most of KdgR is bound by KDG in the mid-exponential phase, which reduces the concentration of free KdgR below the levels required for the binding of both promoters, and fully activates their expression (Fig. 3C). Altogether, the very different induction rates of the two genes thus result from their different basal activity in absence of inducer.

Activation by CRP is also different at the two promoters, but the binding affinities differ only by a factor of 2 (Table 1 and Fig. 3D), resulting in profiles with a similar shape characterized by a sharp peak at transition to stationary phase; however, this peak has a much stronger magnitude for *pelD* due to the three-fold higher activation factor of CRP compared to *pelE* (15).

Altogether, the combined regulation by KdgR and CRP induces both genes almost simultaneously, as observed, short before transition, *i*.*e*., when induction by KDG is still significant while the exhaustion of glucose has already triggered the induction by cAMP. The reader may note a small discrepancy in the predicted expression of *pelE*, which occurs 1-2 hours too late compared to experimental data. Such a deviation was not unexpected though, since *pel* promoters are sensitive to many regulators disregarded in our simplified model (8), as said above. In particular, we note that *pelE* is one of the *D. dadantii* promoters most strongly repressed by DNA supercoiling relaxation (18, 28), which occurs precisely at the transition to stationary phase (29) and is thus expected to displace the *pelE* expression peak toward earlier time, as observed.

## Discussion

### Regulation of *pel* gene expression during plant infection

The presented model quantitatively reproduces the expression dynamics of *pel* genes and pectin degradation, in a culture medium composed of a combination of glucose and pectin. One of the most surprising features of our observations is that most Pel enzymes are produced *after* pectin has been entirely degraded, and they remain therefore unused. This behavior might be rationalized, if we consider the differences between the growth conditions in our stable culture and in the course of plant infection, which is the main context of evolutionary selection.

The availability of carbon sources in this process is summarized in Fig. 4. After invading the plant, *e*.*g*., through a wound, the bacterium experiences a long asymptomatic stage of colonization of the apoplast (intercellular space) where it mostly grows on simple sugars produced by the plant. The maceration phase starts when the available glucose is exhausted, and pectin from the plant cell walls starts to be degraded, resulting in a sudden and rapid extension of soft rot in the plant tissues associated to a massive production of Pels (8). The efficiency of this drastic transition is critical for the success of the infection, since oligogalacturonates (UGA), the product of Pel enzymatic activity, are the main signal inducing the plant defense reactions. Any production of Pels in significant amounts thus triggers a survival race between the plant and the bacteria; if this event occurs before reaching sufficient bacterial density, or if the production boost is not sufficient, bacterial cells will ultimately be destroyed. Does our model provide insights into the regulation events associated to this intrinsically dynamical process?

**Figure 4.**
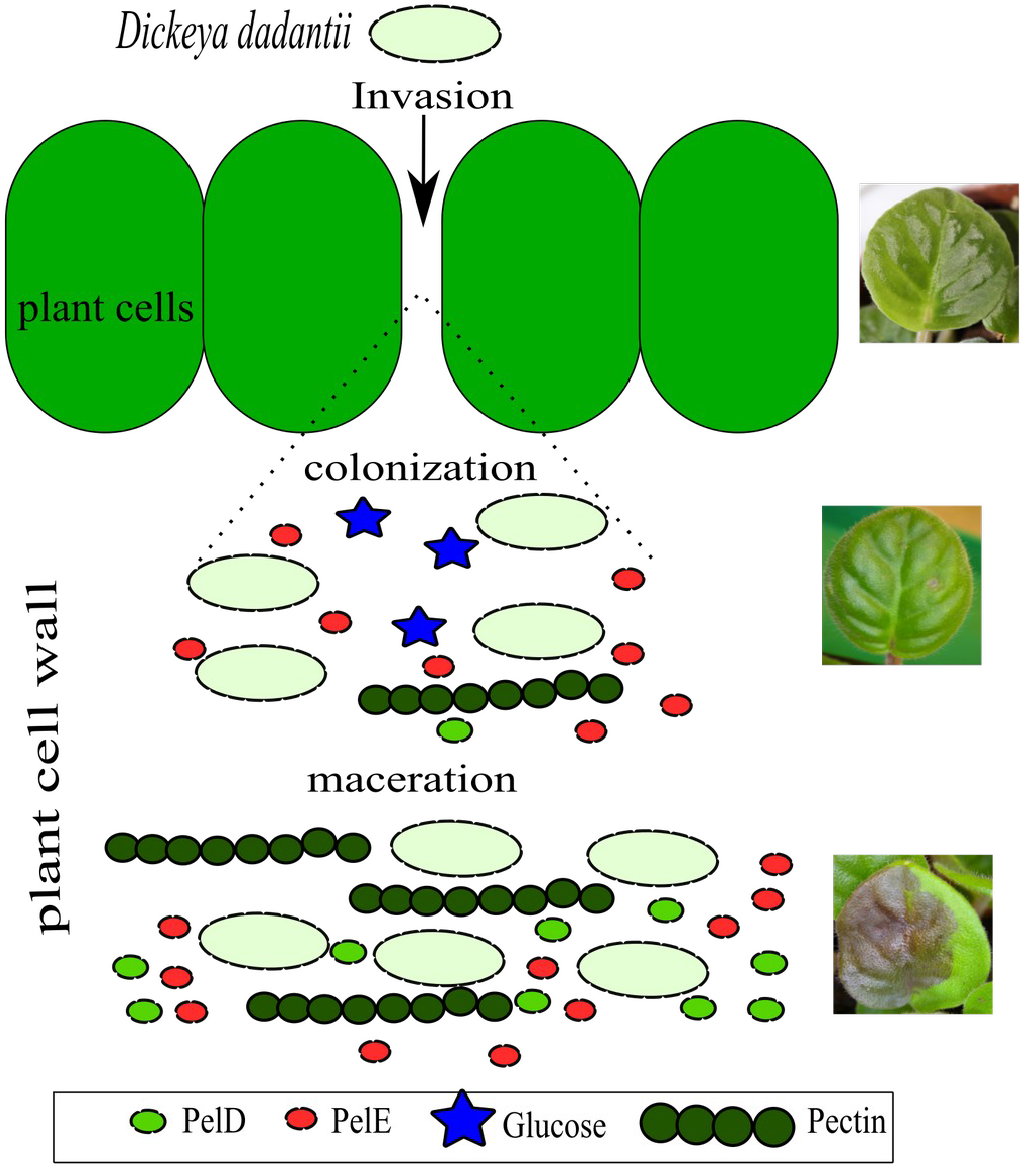
Schematic depiction of the successive steps of plant infection by *D. dadantii*: invasion, colonization, maceration (top to bottom). During colonization (asymptomatic), the bacteria grow on glucose in the plant apoplast. Maceration is characterized by a sudden and massive production of Pels responsible for pectin degradation (visible as soft rot).

Our culture conditions aim at mimicking those encountered during the colonization stage, with bacteria mostly growing on simple sugars, and a level of Pel production remaining sufficiently low to avoid a significant reaction from the plant. Crucially, we have shown that the transition to a high Pel production regime (and thus to the symptomatic phase) is triggered not by an initial event of pectin degradation as suggested earlier (12), but rather by the exhaustion of glucose in the medium (and possibly of other simple sugars in the plant), resulting in a peak of cAMP and *pel* expression boost. The expression of key virulence factors is thus intrinsically related to a metabolic switch of the bacteria.

In the second stage of infection however, the conditions in the plant deviate from those of our stable culture medium. While in the latter, as said above, the low amount of available pectin was already entirely consumed way before the main peak of *pel* expression, in the plant the production of Pels induces a sustained supply of additional pectin as the soft rot propagates in the plant tissues, more similar to the conditions of a continuous bioreactor (30). Based on our modeling, this difference should have a significant regulatory consequence for *pel* genes.

Here, the activating effects of KDG and cAMP were essentially separated in time (6h and 10h, Fig. 3 C and E), with KDG activation having already dropped sharply at the time of *pel* expression (8h). In contrast, with a sustained supply of pectin, KdgR would remain unbound while cAMP peaks, and the simultaneous activation by the two pathways would then induce a considerably stronger expression boost, as required for an efficient infection. Additionally, the different regulatory properties of *pelE* and *pelD* imply that they would be differently affected by this boost. On the one hand, we expect *pelE* to keep playing a crucial role as an inducer of the Pel/KDG feedback loop, because of its higher basal expression level in absence of pectin compared to *pelD* (27). On the other hand, whereas in our data the two genes have a comparable maximal expression level (Fig. 3F), we expect *pelD* to be expressed at a much (approximately three-fold) higher level than *pelE* in the case of a simultaneous activation by both pathways, because it is much more inducible by CRP (Fig. 3E), and should thus play a major role in the massive production of Pels during the maceration phase. In summary, in contrast to our culture conditions where *pelE* plays a predominant role, we expect the two genes to play a qualitatively distinct but equally crucial role in the plant, which is fully consistent with the observation that both *pelD* and *pelE* mutants exhibit phenotypes with strongly reduced virulence (26). The coexistence of these two genes may thus provide a significant evolutionary advantage, possibly explaining their conservation in several *Dickeya* species (*D. dadantii, D. solani, D. zeae, D. dianthicola*) compared to close relatives (*e*.*g*., *D. paradisiaca*) and their ancestor (10).

### Carbon catabolite repression in *Dickeya dadantii*

The competitive uptake of either carbon source presents many similarities, but also interesting differences with the classical example of carbon catabolite repression (CCR) involved in the glucose/lactose switch of *E. coli* (13, 14).

The first apparent and previously identified difference is the monotonous aspect of the growth curve in the mixed medium (Fig. 1A), which contrasts with the diauxic growth of *E. coli* in glucose+lactose (31, 32). Even though pectin is a complex polysaccharide, the consumption of which depends on the production, export and activity of many degrading enzymes, this more continuous pattern previously suggested that pectin consumption starts while glucose is still available, whereas *E. coli* uses lactose only after glucose exhaustion (31). Our KDG quantification data confirm this hypothesis, and even demonstrate that in our culture conditions, pectin is already converted into KDG after 5 hours, *i*.*e*., only half the entire duration of growth. Can we relate this behavior to the underlying regulatory properties of the *pel* genes, and extend the comparison to those of the *lac* operon in *E. coli*?

Fig. 5 gives a schematic depiction of the regulatory systems of these operons controlling the catabolism of the secondary sugar sources in both species (lactose and pectin, respectively). These systems present a conspicuous similarity, with an activation by cAMP-CRP and repression by a pathway-specific regulator, LacI and KdgR respectively, which can be driven away by the degradation product of the sugar, allolactose or KDG respectively. On the other hand, the modes of action of the enzymes involved in the two catabolic pathways are very different, since pectin is a polysaccharide degraded outside the cells by Pels, whereas lactose is a disaccharide and is imported when LacY is present at the cell membrane (32).

**Figure 5.**
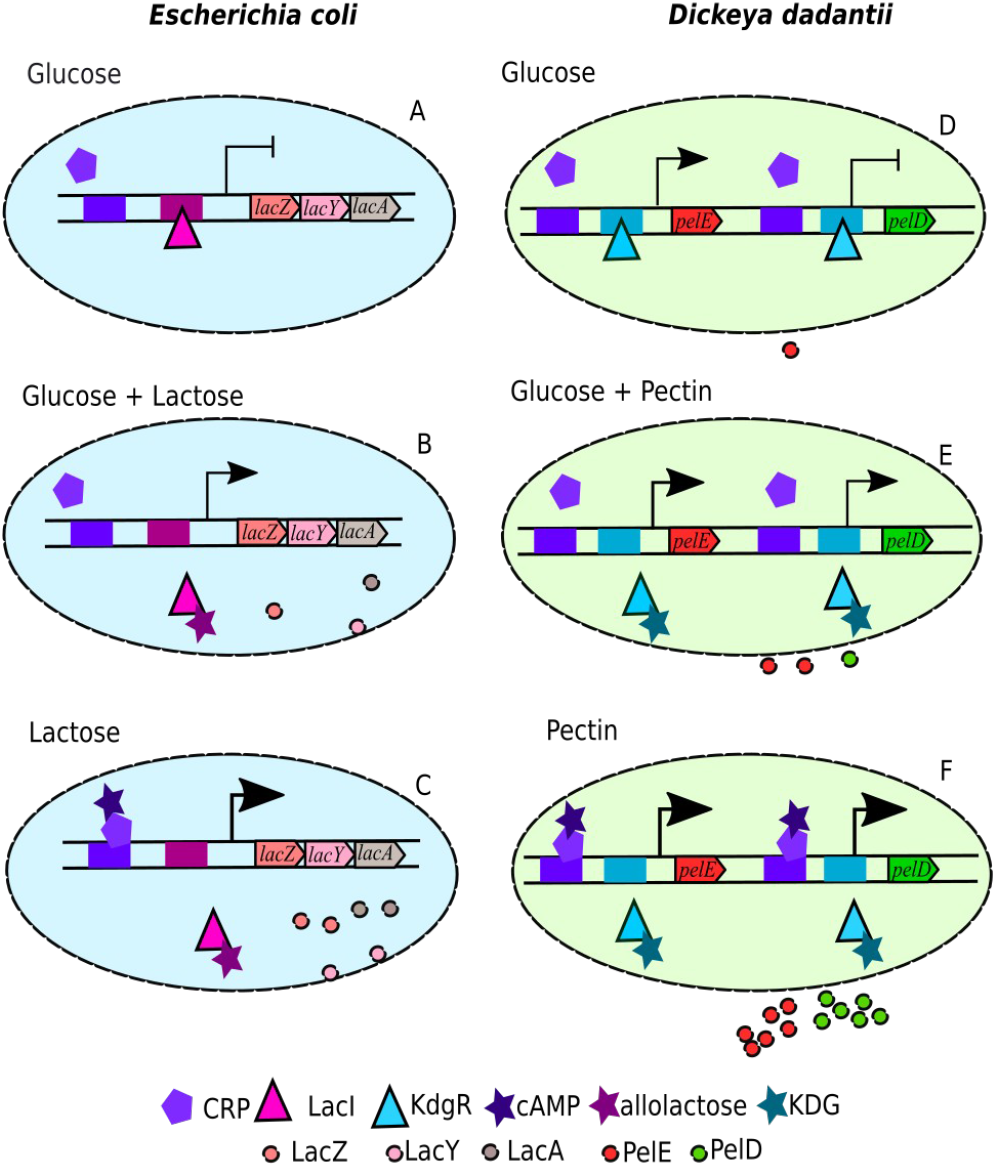
Schematic depiction of the *lac* operon regulation in *E. coli* (left) and *pelED* regulation in *D. dadantii* (right), in presence of glucose, lactose/pectin, or both sources of carbon, illustrating carbon catabolite repression in both systems. Promoters are shown as arrows, with expression levels indicated by arrowhead sizes (and full repression by ⊣). Genes are indicated along the DNA double helix, and regulator binding sites are shown as colored boxes.

In presence of glucose alone (top), the activator CRP is unbound, but the repressor inhibits the expression of *pel*/*lac* genes. In *D. dadantii* however, the presence of a promoter with low repressor affinity (*pelE*) results in a low basal expression and a less drastic repressive effect. When both sources of carbon are present simultaneously (middle), a moderate level of expression can be sustained by the release of the repressor. When only the secondary source is present (bottom), the expression is boosted by the binding of cAMP-CRP.

As visible in Fig. 5, among the two *pel* genes considered in this study, *pelD* is the one with a regulatory system most comparable to the *lac* promoter, exhibiting a strict dependence on the availability of the secondary carbon source. In this figure showing a stationary behavior in each medium, the difference between the two *pel* genes appears quite limited however, and their coexistence would probably not constitute a key advantage if the bacterium lived in a stable or slowly changing environment, as non-pathogenic *E. coli* does most of the time. On the other hand, when both sources of carbon are present (middle panel), the depicted partial expression requires an initial expression event, in order to start producing allolactose/ KDG and sustain subsequent transcription. In *E. coli*, because of the absence of a significant basal level of *lac* expression, this initiation step may be very long, partly explaining why glucose is entirely consumed first (diauxic growth). In *D. dadantii*, the basal expression of *pelE* drastically reduces this initiation time, facilitating the switch in carbon source and resulting in continuous growth. Interestingly, measurements in a *pelE* mutant strain indeed exhibited a severe delay in the expression of *pel* genes, due to a much longer initiation step of the pectin degradation feedback loop (10). As discussed in the previous section, the evolutionary advantage of having coexisting *pelD* and *pelE* genes is thus probably intimately related to the dynamical requirements of *pel* expression in the context of pathogenesis. This expression is part of a race with the host’s defense systems where time plays a prominent role compared to the requirements of non-pathogenic bacteria which are more specifically selected for resource optimization, and where the existence of two *pelE/D* genes with specialized roles (rapidity of the switch vs expression boost) might be key to the success of *D. dadantii*.

## Materials and Methods

### Bacterial Strain and Culture Conditions

*D. dadantii* strain 3937 was used for all experiments described. The cultures were grown at 30 °C in M63 minimal salt medium (33) supplemented with 0.2% (w/v) glucose as primary carbon source and 4% (w/v) PGA for pectin degradation studies. Liquid cultures were grown in a shaking incubator (125 r.p.m.).

### Gene activity assays

*pelE* and *pelD* expression were measured by qRT-PCR as described in (12). Pectate lyase enzymatic activity assays were performed on toluenized cell extracts in 1 mL cell culture harvested at predetermined time points along the growth of the bacteria. The degradation of PGA by pectate lyases is monitored by absorption spectrometry of unsaturated oligogalacturonides at 230 nm (34). Pel activity is expressed as µmol of unsaturated products liberated per minute and per ml of enzymatic extract, as described in (12). All experiments were carried with two biological replicates.

### Bacterial samples preparation for KDG and cAMP quantification

The bacterial culture samples were harvested at predetermined time points after measuring the optical density (OD), centrifuged at 8000 rpm for 6 minutes to separate cells from supernatant medium. Bacterial pellet was freeze dried with liquid nitrogen for storing. Cells and medium samples were stored separately at -80°C. In order to process for quantification the cell pellet was resuspended in 10 ml cold solution of 2.5 % (v/v) TFA, 50% (v/v) acetonitrile. The dissolved lysed pellet was maintained at -80°C for 15 minutes and thawed at room temperature. The solution was then centrifuged at 20.000*g* for 5 minutes. The clear supernatant containing the metabolites (10 ml) was separated from the pellet and then lyophilized overnight. The dry powder was dissolved in 1 ml of 0.1 M HCl and centrifuged again to remove particles. Culture medium is directly made up to .1 M HCl with 1 M HCl. The measurements were carried in two biological replicates in glucose+PGA medium (and one in the glucose control medium).

### KDG and cAMP quantification using HPLC

KDG metabolite was quantified using a novel HPLC technique developed inhouse as described in (17). The carbonyl group was derivatized with o-phenylenediamine and eluted with a buffer consisting of 0.005% trifluoroacetic acid and 60% acetonitrile.

cAMP metabolite was quantified by HPLC after derivatization with 2-chloroacetaldehyde (21) (Supplementary Figure S2). Bacterial cell and extracellular medium samples extracts are suspended in 0.1 M HCl. Equal amounts of sample in 0.1 M HCl and 1 M sodium acetate are mixed and 10% v/v of 2-chloroacetaldehyde solution is added to the mixture and incubated at 80°C for 30 min with continuous shaking (35). The appropriate amounts of incubated sample are injected onto an Uptisphere C18 column (Interchim) for separation of the purine derivative adducts. Buffer A was composed of 0.005 % trifluoroacetic acid (TFA) in water and Buffer B of 100% methanol. The flow rate was set-up at 35 °C, and run at 1 ml/min using the following gradient: initial 90% A, 10% B; linear gradient from 0 to 20 min up to 70% A, 30% B; 22 min 100% B; 25 min 25% B then re-equilibration to initial condition from 28 to 30 min (see the profile in Supplementary Figure S3). The mean retention time for cAMP was between 13.04 to 13.10 min. The associated peak is well-separated from neighbor peaks characteristic of other compounds of similar structure (Supplementary Figure S3B), and can be further distinguished by its characteristic absorption/emission spectrum identical to that of the purified molecule (data not shown). The quantification is performed from a standard curve (Supplementary Figure S4). Raw extracellular concentration values are shown in Supplementary Figures S5 (glucose+PGA medium) and S6 (glucose medium). The model used to infer intracellular cAMP concentrations is adapted from (22) and described in Supplementary Information.

### Model

The system dynamics is simulated with a set of deterministic ordinary differential equations relating the concentrations of all considered species (see Fig. 2 and Supplementary Information). Bacterial growth follows a logistic equation where the maximal growth rate and final density depend linearly on the available nutrients (sucrose and/or PGA) (12). All binding reactions are described as thermodynamic equilibria, in particular those between the metabolites (KDG and cAMP) and the regulatory proteins (KdgR and CRP dimers, respectively), and between the latter and the two *pel* promoters. Promoter activities follow a thermodynamic model of transcription (25) involving the activator cAMP-CRP and repressor KdgR (Table 1). All enzymatic reactions are treated with Michaelis-Menten kinetics. The list of model parameters is provided in Supplementary Table S2. All thermodynamic and regulatory parameters were obtained experimentally (12, 15, 19) as well as most kinetic parameters. The 5 remaining parameters (Supplementary Table S3) were numerically estimated by fitting a set of observed quantities (*pel* expression peak time and ratio in presence and absence of PGA, KDG concentration peak time and magnitude) using the truncated Newton method. The dynamical system was simulated with the Euler algorithm, using a constant timestep of 0.3 min ensuring numerical stability. All data analyses and computations were carried in Python using NumPy, Scipy and Pandas libraries.

## Acknowledgments

We thank the whole CRP team for fruitful discussions, and Hans Geiselmann for suggestions and critical reading of the manuscript.

## Funding

The authors acknowledge INSA Lyon for a Bonus Qualité Recherche 2016 grant, and a Bourse Enjeux 2017 fellowship to S. Martis.

**Figure 1. (A)** Growth curves of *D. dadantii* in M63 minimal medium supplemented with glucose (gray) or glucose+PGA (black). **(B)** Time course of PGA degradation (dashed line, data from (12)) and intracellular KDG concentration measured by HPLC (solid line). The horizontal dashed line (0.4 mM) indicates the KDG-KdgR affinity: above this level, repression is relieved and *pel* expression is strongly induced. **(C)** Time course of measured extracellular cAMP concentration (dashed line) and inferred intracellular cAMP concentration (solid line, with shaded confidence area), for cells grown in glucose (gray) or glucose+PGA (black). The dashed horizontal line (10 µM) indicates the cAMP-CRP binding affinity (16): above this level, *pel* expression is activated. Raw data points are shown in Supplementary Figures S5 and S6. **(D)** *pelE* expression measured by qRT-PCR (dashed line) and total Pel enzymatic activity in glucose (gray) or glucose+PGA (black) media.

**Figure 2**. Diagram of the components and reactions considered in the model. The two metabolites, cAMP and KDG, indirectly activate the expression of *pel* genes by binding their associated regulators: cAMP allows CRP to bind and activate the promoters, whereas KDG relieves KdgR repression, respectively. Pel enzymes are instantaneously exported and degrade PGA (pectin) into UGA, which is imported and converted into KDG, triggering a positive feedback loop. KdgR is synthesized at a constant rate.

**Figure 3**. Modeling of *pel* regulatory pathways and expression in bacteria grown in minimal medium supplemented with glucose+PGA (left column, A-G) or glucose (right, H-N). Model results are shown as solid lines, and experimental data as dots. The transition time is shown as a dashed gray vertical line. **(A)** KDG intracellular concentration superimposed with HPLC direct measurement data exhibiting a peak after around 6 hours of growth; **(B)** concentration of intracellular free KdgR dimer, with horizontal dashed lines showing the binding affinities of the dimer for the *pelE* (red) and *pelD* (green) promoters, indicating the thresholds of repression; **(C)** *pelE* and *pelD* (depicted in red and green respectively throughout the figure) activation curves based on KdgR binding, exhibiting a de-repression in the middle of exponential phase due to high intracellular levels of KDG; **(D)** intracellular concentration of the cAMP-CRP complex inferred from HPLC measurement of cAMP in the medium (Fig. 1B); the gray area is the 95% confidence interval due to experimental variations, and dashed horizontal lines indicate the binding affinities for *pelD/E* promoters; **(E)** *pel* activation curves based on cAMP-CRP binding, with a boost occuring at transition; **(F)** *pel* expression time course based on combined regulation by KdgR and CRP, superimposed with qRT-PCR datapoints peaking around 8h; **(G)** extracellular concentration of Pel enzymes, either from individual genes (red and green) or the combination of both (black), superimposed with measurements of enzymatic activity (dots) reflecting total Pel enzymes, including those not considered in our modeling. **(H-N)** Same legends as A-G: in absence of PGA, the dilution of KdgR during growth is sufficient to partly relieve *pelE* repression, while cAMP induces expression of both *pels* close to transition. The timing and expression levels of both genes is accurately reproduced by the model in both conditions, as well as the inducing effect of PGA.

**Figure 4**. Schematic depiction of the successive steps of plant infection by *D. dadantii*: invasion, colonization, maceration (top to bottom). During colonization (asymptomatic), the bacteria grow on glucose in the plant apoplast. Maceration is characterized by a sudden and massive production of Pels responsible for pectin degradation (visible as soft rot).

**Figure 5**. Schematic depiction of the *lac* operon regulation in *E. coli* (left) and *pelED* regulation in *D. dadantii* (right), in presence of glucose, lactose/pectin, or both sources of carbon, illustrating carbon catabolite repression in both systems. Promoters are shown as arrows, with expression levels indicated by arrowhead sizes (and full repression by ⊣). Genes are indicated along the DNA double helix, and regulator binding sites are shown as colored boxes.

## List of Supplementary Data

**Supplementary Table S1**. List of variables in the dynamical system

**Supplementary Table S2**. Parameters of the model with fixed value

**Supplementary Table S3**. Unknown parameters adjusted on the data

**Supplementary Table S4**. Growth parameters

**Supplementary Figure S1**. Detailed pectin degradation pathway

**Supplementary Figure S2**. Derivatization of cAMP with 2-chloroacetaldehyde

**Supplementary Figure S3**. Elution profile of cAMP by fluorescence detection at 418 nm after specific excitation at 278 nm, in a sample containing purified cAMP (A) or extracellular culture extracts in M63 Glucose+PGA medium (B). The fluorescence signal is shown in red (left scale), and the proportion of methanol (vs trifluoroacetic acid) in the elution buffer is shown as a dashed black line (right scale).

**Supplementary Figure S4**. Standard curve of cAMP titration.

**Supplementary Figure S5**. Raw datapoints of extracellular cAMP concentrations in M63 Glucose+PGA medium.

**Supplementary Figure S6**. Raw data points of extracellular cAMP quantifications in M63 Glucose medium

